# Temporal analysis suggests a reciprocal relationship between 3D chromatin structure and transcription

**DOI:** 10.1101/2022.05.05.490836

**Authors:** Kathleen S. M. Reed, Eric S. Davis, Marielle L. Bond, Alan Cabrera, Eliza Thulson, I. Yoseli Quiroga, Shannon Cassel, Kamisha T. Woolery, Isaac Hilton, Hyejung Won, Michael I. Love, Douglas H. Phanstiel

## Abstract

To infer potential causal relationships between 3D chromatin structure, enhancers, and gene transcription, we mapped each feature in a genome-wide fashion across eight narrowly-spaced timepoints of macrophage activation. Enhancers and genes connected by loops exhibited stronger correlations between histone H3K27 acetylation and expression than can be explained by genomic distance or physical proximity alone. Changes in acetylation at looped distal enhancers preceded changes in gene expression. Changes in gene expression exhibit a directional bias at differential loop anchors; gained loops are associated with increased expression of genes oriented away from the center of the loop, while lost loops were often accompanied by high levels of transcription with the loop boundaries themselves. Taken together, these results are consistent with a reciprocal relationship in which loops can facilitate increased transcription by connecting promoters to distal enhancers while high levels of transcription can impede loop formation.

**HIGHLIGHTS:**

- LPS + IFNγ triggers genome-wide changes in chromatin looping, enhancer acetylation, and gene expression
- Looped enhancer-promoter pairs exhibit ordered and correlated changes in acetylation and expression
- Changes in gene expression exhibit a directional bias at differential loop anchors
- Lost loops are associated with high levels of transcription within loop boundaries

## INTRODUCTION

3D chromatin structure is thought to play a critical role in gene expression, cellular identity, and organismal development by modulating contact frequencies between gene promoters and distal regulatory elements such as enhancers^1^. Alterations in 3D chromatin architecture have been associated with developmental abnormalities and human disease^2–6^. Despite growing knowledge regarding the proteins and molecules that govern 3D chromatin architecture, the relationship between 3D chromatin architecture and gene transcription is less certain. While some functional connections between chromatin interactions and transcription have been established, the degree to which 3D chromatin structure shapes—or is shaped by—transcription remains unclear.

The continued development of chromatin conformation capture (3C) based technologies has provided valuable insights into the mechanisms driving 3D chromatin structure^7–13^. In particular, genome-wide approaches including Hi-C have revealed tens of thousands of loops throughout the human genome, many of which connect regulatory elements such as enhancers to gene promoters. With some notable exceptions^2,14,15^, the majority of loops are bound at each anchor by CTCF and are formed via loop extrusion by the cohesin complex^16,17^. Mapping these loops across cell types and biological conditions has revealed cell-type-specific looping events that often correlate with differences in gene transcription^9,18–20^.

Despite these advances, the mechanisms and degree to which looping drives transcriptional changes is far less certain. A widely held hypothesis is that chromatin loops facilitate transcriptional activation by increasing the frequency of interactions between enhancers and gene promoters; however, studies that removed looping genome-wide have produced conflicting results. Acute depletion of cohesin in a human cancer cell line was sufficient to eliminate cohesin-bound loops but had only a modest effect on transcription, casting doubt on the importance of DNA looping for transcriptional control^21^. In contrast, deletion of the cohesin loading factor, NIPBL, in mouse liver cells in vivo induced transcriptional changes of thousands of genes^22^. In addition, deletion of cohesin significantly impacted the ability of mouse macrophages to mount a proper transcriptional response to a microbial stimulus^23^, which suggests that loops might be specifically important for regulating changes to (as opposed to maintenance of) transcriptional signatures.

Mounting evidence also suggests that transcription can shape 3D chromatin structure, although the exact relationship remains unclear. Several studies have shown that transcription can displace cohesin and condensin complexes^24–26^. For example, knocking down CTCF and the cohesin unloader WAPL caused cohesin to accumulate at the 3’ end of highly transcribed genes, suggesting that cohesin may be relocated by transcription in the absence of boundary elements^24^. At least two studies have shown that transcription-induced displacement of SMC complexes results in altered chromatin structure. Macrophages infected with influenza A, which inhibits transcription termination, revealed readthrough transcription that displaced CTCF and repositioned cohesin at the 3’ end of genes, disrupting existing chromatin structure^27^. Fibroblasts undergoing senescence exhibit de novo transcription-dependent cohesin peaks at the 3’ end of select genes, resulting in newly formed loops^28^. These findings are supported by in vitro experiments performed on DNA “curtains” showing that RNA polymerase or other translocases can push cohesin; however, more recent studies that suggest molecules as large as 200 nm may be able to pass through SMC complexes^29–31^.

One approach to dissect causal relationships between looping and transcription—while circumventing genome-wide perturbations with potential knock-on effects—is to quantify changes in looping, transcription, and other regulatory features across biological timecourses. Indeed, 3C-based timecourses of biological transitions have produced valuable insights into the dynamics of 3D chromatin architecture^32–39^. For example, D’Ippolito *et al.* characterized differential looping at 4 timepoints following glucocorticoid treatment and found that on average loops changed maximally at 4 hours whereas gene expression changed maximally at 9 hours^20^. This timing is consistent with a regulatory relationship, though the relatively broad spacing of timepoints made temporal ordering of individual pairs of loops and genes more difficult. In another study, Beagan *et al.* used 5C to identify differential looping events in activated neurons in time frames as short as 20 minutes^40^; however, these studies focused on just a handful of genomic loci.

To characterize the temporal order of regulatory events and infer potential causal relationships, we mapped 3D chromatin architecture, histone H3 K27 acetylation, chromatin accessibility, and gene expression across eight timepoints of macrophage activation. Narrowly spaced timepoints allowed correlation and temporal ordering of events at a locus by locus level. These analyses provided insights into the putative causal relationships between these events which were consistent with a reciprocal relationship between chromatin looping and gene transcription.

## RESULTS

### LPS + IFNγ triggers genome-wide changes in chromatin looping, enhancer acetylation, and gene expression

To understand how chromatin loops and enhancers work together to regulate gene transcription in response to external stimuli we conducted an eight-point timecourse of human macrophage activation (**Fig 1A**). Human macrophages derived from the THP-1 monocytic cell line were stimulated with 10 ng/mL lipopolysaccharide (LPS) and 20 ng/mL interferon-gamma (IFNγ) and collected at eight timepoints (0, 0.5, 1, 1.5, 2, 4, 6, and 24 hours). At each timepoint we profiled 3D chromatin structure using in situ Hi-C^9^, putative enhancer activity using ChIP-seq targeting histone H3K27 acetylation, chromatin accessibility using ATAC-seq^41^, and gene expression using RNA-seq.

**Figure 1.**
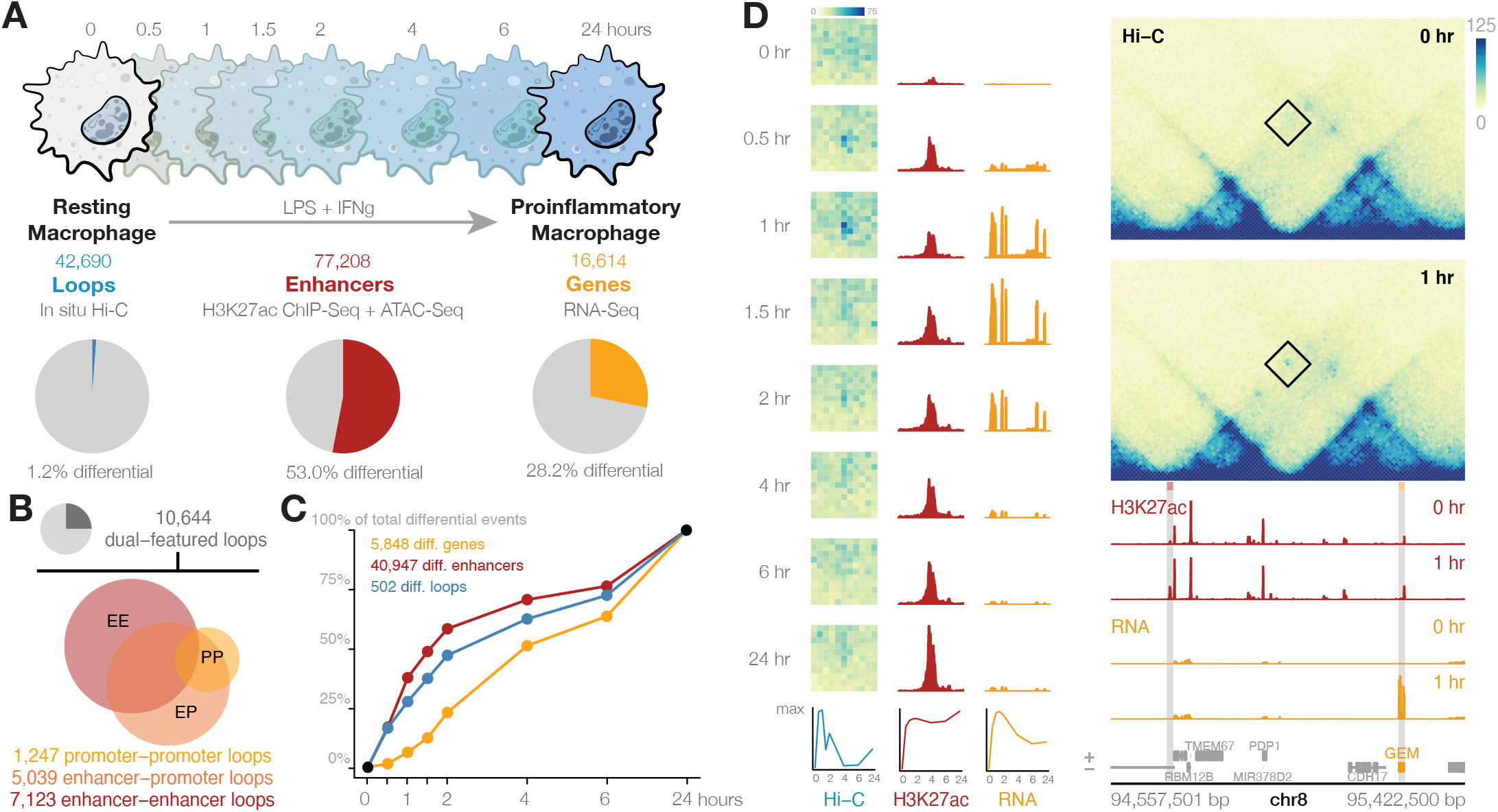
Multi-omic timecourse of macrophage activation physically and temporally connects regulatory events. **(A)** Experimental design to identify changes to 3D chromatin structure, enhancers, and gene expression across eight timepoints during macrophage proinflammatory activation. **(B)** Fraction and number of loops that connect two distal elements. **(C)** A cumulative sum of differential events identified by each timepoint reveals the relative timing of changes to genes, loops, and enhancers. **(D)** Intersecting differential chromatin loops, enhancers, and genes provides the regulatory context of transcriptional changes. At this region, a 570-Kb loop connects the promoter of the GEM gene to a distal enhancer. The enhancer’s activity peaks 30 minutes before gene expression but remains high throughout the treatment, while the loop connecting them fades alongside gene expression after 2 hours.

With eight timepoints and roughly 2 billion contacts per Hi-C map, this represents one of the most comprehensive characterizations of 3D chromatin changes to date^20,42^. To comprehensively catalog long-distance chromatin interactions, we further combined our maps from each timepoint into a single, ultra-deep “Mega” map comprising 24.5 billion reads and 15.6 billion chromatin contacts **(Fig S1A-B)**. This increased read depth provided the power to identify over 10,000 additional loops that were undetectable at the resolution of individual time-points **(Fig S1C)**. Loops from each timepoint as well as the Mega map were then merged, combining any loops with both anchors within 20 kb, to provide 42,690 total loops for this study.

To identify potential regulatory connections among these loops, we classified putative enhancers (henceforth called enhancers) as loci with overlapping ATAC-seq and histone H3K27 acetylation peaks that did not overlap gene promoters (see methods). Intersecting these enhancers with chromatin loops revealed 5,039 enhancer-promoter loops (**Fig 1B**). The regulatory activity of enhancers was inferred via quantification of histone H3K27ac at each enhancer. Finally, we used stranded rRNA-depleted RNA-seq at each timepoint to quantify the potential impacts of these loops and enhancers on gene expression.

Differential analysis using the DESeq2 package^43^ identified statistically significant genome-wide alterations in DNA looping, enhancer activity, and gene expression at each timepoint (**Fig 1C**, **Table S1-3**). The transcriptional changes we observed are consistent with previously established profiles of inflammatory activation **(Fig S2)**. Only 1.2% (220 up, 282 down) of loops were detected as differential in at least one timepoint, compared to 53.0% (21,858 up, 19,089 down) of enhancers and 28.2% (3,025 up, 2,823 down) of genes. Of these 502 differential loops, 79 were detected only at intermediate timepoints and were not visible at either 0 or 24 hours, highlighting the insights offered from this level of temporal resolution. On average enhancers and loops changed faster than genes, with 58.4% of differential enhancers and 47.2% of differential loops changing significantly within the first 2 hours of LPS + IFNγ treatment compared to only 23.1% of genes (**Fig 1C**). This temporal lag between changes in loops and enhancers compared to changes in gene expression is consistent with our understanding of loops and enhancers as regulators of gene transcription, and highlights the power of using temporal analysis to generate hypotheses about causal relationships^44–47^.

Integrating the resulting multi-omic data provided insights into gene regulatory mechanisms of macrophage activation. An example of this concept can be seen at the *GEM* locus on chromosome 1 (**Fig 1D**). The *GEM* gene is transiently upregulated during LPS + IFNγ treatment, with expression peaking between 1 and 2 hours. An enhancer 570 Kb downstream of the *GEM* promoter becomes acetylated and physically looped to the promoter of *GEM* after only 30 minutes of treatment. While acetylation of this distal enhancer remains high throughout the treatment, the contact frequency of this regulatory loop changes along the same temporal pattern as *GEM* expression, albeit preceding transcriptional changes by approximately 30 minutes. Taken together, these data are supportive of a model in which 3D contacts between an active enhancer and gene promoter play a causal role in transcriptional changes. Throughout the rest of this paper, we explore these relationships quantitatively on a genome-wide scale.

### Looped enhancer-promoter pairs exhibit ordered and correlated changes in acetylation and expression

The importance of chromatin looping for transcriptional regulation remains unclear, as studies disrupting chromatin loops comprehensively throughout the genome have produced mixed results^21–23^. Ablation of loops in the human colorectal cancer cell line HCT-116 only altered the expression of a handful of genes^21^. In contrast, loss of loops in murine liver cells and macrophages responding to LPS induced thousands of transcriptional changes^22,23^. Differences in biological systems, cellular contexts, and even the method of loop disruption could all potentially explain the conflicting findings.

We investigated our data to see if it supported a role for looping in gene regulation in response to external stimuli. Our analyses were based on the assumption that if loops play a role in transcriptional control, enhancer-gene pairs should exhibit correlated changes in histone H3K27ac and gene expression. Because only a small fraction of loops change over time, all loops were used to connect enhancers to promoters regardless of differential status. In total, this involved 5,039 enhancer-promoter loops featuring 4,093 unique genes, 1,483 of which were differential. In total, 25.4% of differential genes were connected to a distal enhancer via a chromatin loop. We investigated the temporal patterns of these looped enhancer-promoter pairs and compared them to sets of enhancer-promoter pairs that were matched for either genomic distance or contact frequency using the matchRanges function available from the nullranges R/ Bioconductor package (**Fig 2A-B**).

**Figure 2.**
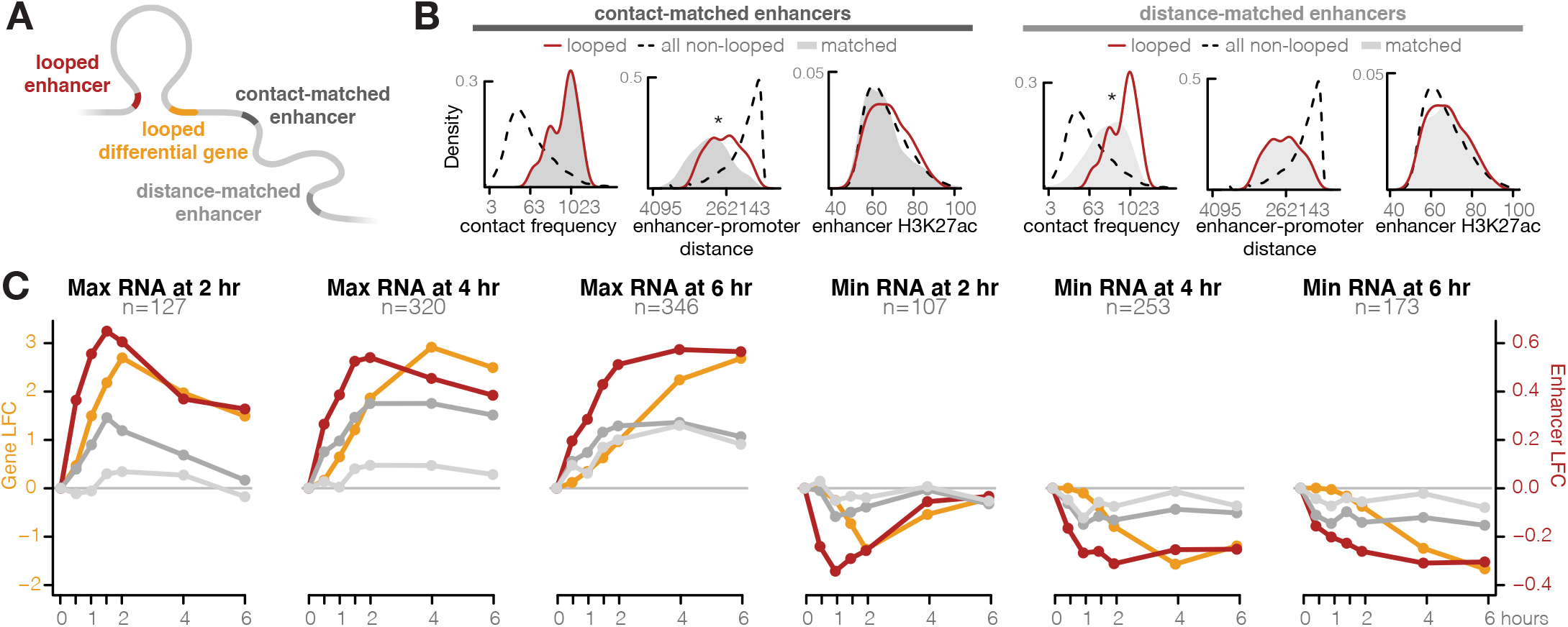
Enhancer acetylation and gene expression correlate most highly at looped enhancer-promoter pairs. **(A)** Distal enhancers looped to the promoters of differential genes were compared to matched enhancers of equal H3K27ac and contact frequency (dark grey), or distance (light grey). **(B)** Representative distributions of contact- and distance-matched enhancers compared to the pool of non-looped enhancer-promoter pairs and the looped subset. Compared to looped pairs, contact-matched enhancers are closer on average in base pairs (Wilcoxon rank-sum test, p-value < 10-6), while distance-matched enhancers are in less frequent contact (Wilcoxon rank-sum test, p-value < 10-11). Both sets of matched enhancers have similar H3K27ac levels to the looped pairs (Wilcoxon rank-sum test, p-value .028, .86). **(C)** Average log2 fold-change of gene expression (gold) for genes reaching minimum or maximum fold-change at 2, 4, or 6 hours are compared to log2 fold-change of their looped enhancers (red), contact-matched enhancers (dark grey), and distance-matched enhancers (light grey). Looped enhancers correlate significantly with changes in gene expression, to a larger extent than matched enhancers. Contact-matched enhancers tend to correlate better than distance-matched enhancers at upregulated genes. Changes in distal enhancer H3K27 acetylation precede changes in gene expression among all timescales, and among both up- and downregulated genes.

To explore the correlation of enhancers and genes over time, we clustered our differential genes based on the timepoint at which they exhibited their maximal up- or down-regulation with respect to the 0 hour timepoint and plotted their average normalized expression (**Fig 2C, yellow lines**). Only clusters peaking at intermediate timepoints and with more than 100 genes are shown. For each gene cluster, we identified enhancers that were connected to those genes via a chromatin loop and plotted their average normalized histone H3K27ac signal (**Fig 2C, red lines**). All 6 clusters revealed a clear correlation between histone H3K27ac and gene expression at looped enhancer-promoter pairs supporting the idea that looped pairs are functionally connected. Interestingly, the changes in acetylation preceded changes in gene expression by 30-60 minutes. This lag is consistent with enhancer activation causing changes to gene expression. However, chromatin loops occur over relatively short distances (median ~390Kb) and at such short distances, even non-looped enhancers and promoters exhibit elevated chromatin contact frequencies compared to randomly selected enhancers and genes across the genome. Therefore, the correlation between looped enhancers and promoters that we observe could be explained by genomic distance alone.

To determine if looped enhancer-promoter pairs exhibited higher correlation than expected given their genomic distance, we compared looped enhancer-promoter pairs to non-looped enhancer-promoter pairs that were matched for genomic distance (**Fig 2A**). As expected, distance-matched non-looped enhancer-promoter pairs were characterized by a lower contact frequency than looped enhancer-promoter pairs. Distance-matched, non-looped, enhancer-promoter pairs exhibited some degree of correlation (**Fig 2B, light grey lines**); however, the correlation was weaker than that observed at looped enhancer-promoter pairs. Thus distance alone does not account for the enhancer-promoter correlations observed at loop anchors and offers further support for the functional role of loops in enhancer-based gene regulation.

One explanation for how loops exhibit transcriptional control is by increasing contact frequencies between enhancers and their target genes. To determine if looped enhancer-promoter pairs exhibited a higher correlation than expected given their contact frequency, we compared looped enhancer-promoter pairs to non-looped enhancer-promoter pairs that were matched for contact frequency. The genomic distance between contact-frequency-matched, non-looped enhancers and promoters was on average far shorter than looped enhancers and promoters (**Fig 2B**). Surprisingly, while contact-matched pairs exhibited a stronger correlation than distance-matched pairs, the correlation was still weaker than that observed at looped enhancer-promoter pairs (**Fig 2C, dark grey lines**). We confirmed these results using data from our previously published study of monocyte differentiation (**Fig S3**)^19^. There too, looped enhancer-promoter pairs exhibited better correlation than enhancer-promoter pairs that were matched for either distance or contact frequency. This was surprising and suggests that the presence of a chromatin loop may facilitate a functional regulatory connection through mechanisms beyond simply increasing their frequency of physical proximity. We explore some possible explanations for this in the discussion.

### Changes in gene expression exhibit a directional bias at differential loop anchors

Given the correlation that we observed between acetylation and gene expression at opposite ends of chromatin loops, we hypothesized that changes in looping would be associated with altered transcription of genes at loop anchors and that the directionality of changes in expression would match that of the changes in looping. To test this, we used k-means clustering to identify four categories of differential looping: gained early, gained late, lost early, and lost late. Examples of loops from each cluster are shown in **Fig 3A**. Differential loops spanned approximately 170-200 kb on average, with the exception of gained late loops which were much larger with an average length of 610 kb (**Fig S4A**). Next, we calculated the percentage of genes at each set of loop anchors that were significantly up or downregulated in response to LPS + IFNγ (**Fig 3B, S4B**). Anchor genes were defined by overlapping gene promoters with loop anchors (see methods). Gained loop anchors were enriched for the promoters of upregulated genes (permutation test, n=10,000, p-value < 0.05) which is consistent with findings from previous work by our lab and others that have associated increased looping with increased transcription of anchor genes^9,19^, and generally supports a causal role for looping in transcriptional control. Unintuitively, however, lost loops were also associated with increased transcription of anchor genes (permutation test, n=10,000, p-value < 0.05). Though surprising, this is consistent with data from Rao *et al.,* which showed that removal of DNA loops is not necessarily accompanied by decreased transcription of anchor genes^21^.

**Figure 3.**
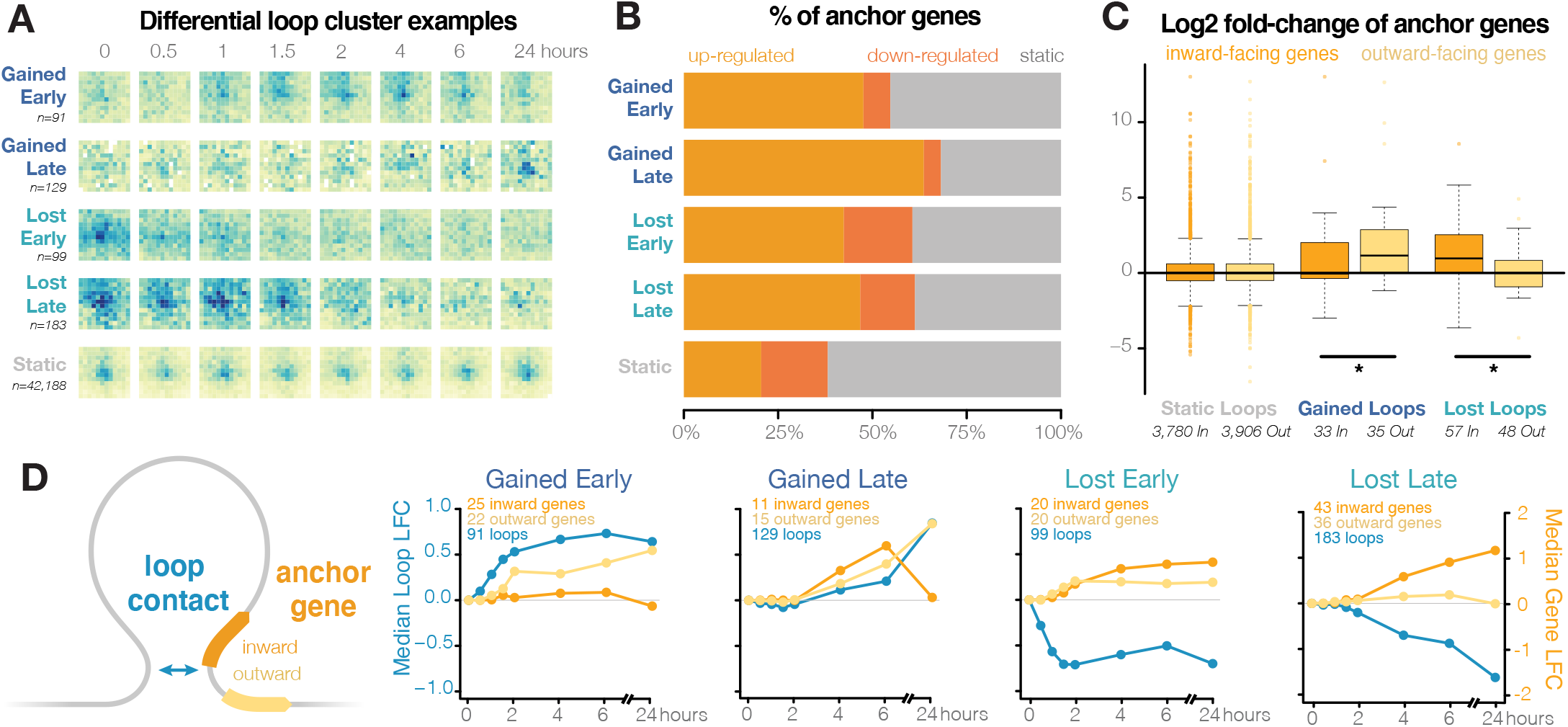
Upregulated genes anchored at differential loops exhibit directionality bias. **(A)** 502 differential loops were clustered by their timing and direction. Representative loops are shown for each cluster. **(B)** The anchors in all differential loop clusters are enriched for upregulated genes. **(C)** Distributions of log2 fold-changes of unique genes with promoters in the anchors of static and differential loops. Anchor genes were classified by whether they are oriented towards (inward, orange) or away from (outward, yellow) the center of the loop. Among genes at gained loop anchors, the fold-change of outward-facing genes is significantly higher than inward-facing genes, while the opposite trend is seen among genes at lost loops (Wilcoxon rank-sum test, p-value < 0.05). **(D)** Average log2 fold-change of differential loops (blue) and inward - and outward-facing genes (orange, yellow) with promoters overlapping those loop anchors.

To explore this further, we separately analyzed anchor gene expression based on whether genes were oriented towards or away from the center of the loop. Intriguingly, genes at the anchors of gained and lost loop classes exhibited different directional biases (**Fig 3C, S4C**). At gained loops, outward-oriented anchor genes exhibited significantly more increased expression than inward-oriented genes (Wilcoxon rank-sum test, p-value < 0.05). In contrast, at lost loops, inward-oriented anchor genes exhibited more increased expression than outward-oriented genes (Wilcoxon rank-sum test, p-value < 0.05). Similar trends can be seen using differential loops from monocyte-to-macrophage differentiation (**Fig S4D**). To investigate this further, we examined the temporal profiles of differential loops and anchor genes. For each loop cluster, we calculated the average fold-change of inward- and outward-facing anchor genes (**Fig 3D**). At gained loops, contact frequency and transcription of anchor genes were positively correlated over time, particularly for loops oriented away from the center of the loop. The gained early loops exhibited increased contact frequency 30-60 minutes prior to the increased transcription of outward-oriented anchor genes which is consistent with the notion of loops playing a causal role in gene expression. The gained late loops showed correlated changes in outward-oriented anchor gene expression but because they changed most drastically between 6 and 24 hours, the timepoints were not close enough to observe a temporal lag. In contrast, at lost loops, contact frequency and transcription of anchor genes were inversely correlated, particularly for loops oriented towards the center of the loop.

### Lost loops are associated with high levels of transcription within loop boundaries

One possible explanation for the directional biases we observe at differential loop anchors is that transcription may be antagonistic to loop extrusion and that high levels of transcription at loop anchors, or within the loop itself, may destabilize loop extrusion complexes. This would agree with several previous studies highlighting the ability of RNA polymerase to push and/or displace cohesin^24,27^.

To determine if antagonism between transcription and loop extrusion could explain the increased expression we observed at lost loop anchors, we explored the absolute and relative levels of transcription occurring within the boundaries of differential loops. Since the majority of transcription occurs at introns, which are generally not captured in our RNA-seq data, we devised an inferred transcription score (ITS) to roughly estimate the levels of transcription for every 10 Kb bin in the genome using our RNA-seq data (see methods). Briefly, the transcript per million (TPM) value for each gene was assigned to every genomic bin covered by the gene body. Values were summed for bins that overlapped multiple genes.

Using our inferred transcription scores, we observed that gained loops have relatively low levels of internal transcription at all timepoints (average ITS ≤ 50, **Fig 4A**). In contrast, decreasing loops achieved much higher average levels of internal transcription during the timecourse (**Fig 4A,** ITS > 50 at most timepoints), and the amount of internal transcription is inversely correlated with changes in loop strength (R^2^ is −0.59 for lost early and −0.99 for lost late loops). To determine how big of a change in ITS was required for a decrease in loop strength, we explored how the changes in ITS within a loop correlated with loop fold change. Loops with a mean increase ITS of 10 or more exhibited a statistically significant decrease in loop strength (Wilcoxon signed rank test, p-value < 0.01 **Fig 4B**). Visualizing transcription relative to the loop boundaries **(Fig 4C)** further confirmed these findings. Transcription was enriched outside of the anchors of gained loops but between the anchors of lost loops. These data are consistent with a model in which high levels of transcription antagonize loop extrusion.

**Figure 4.**
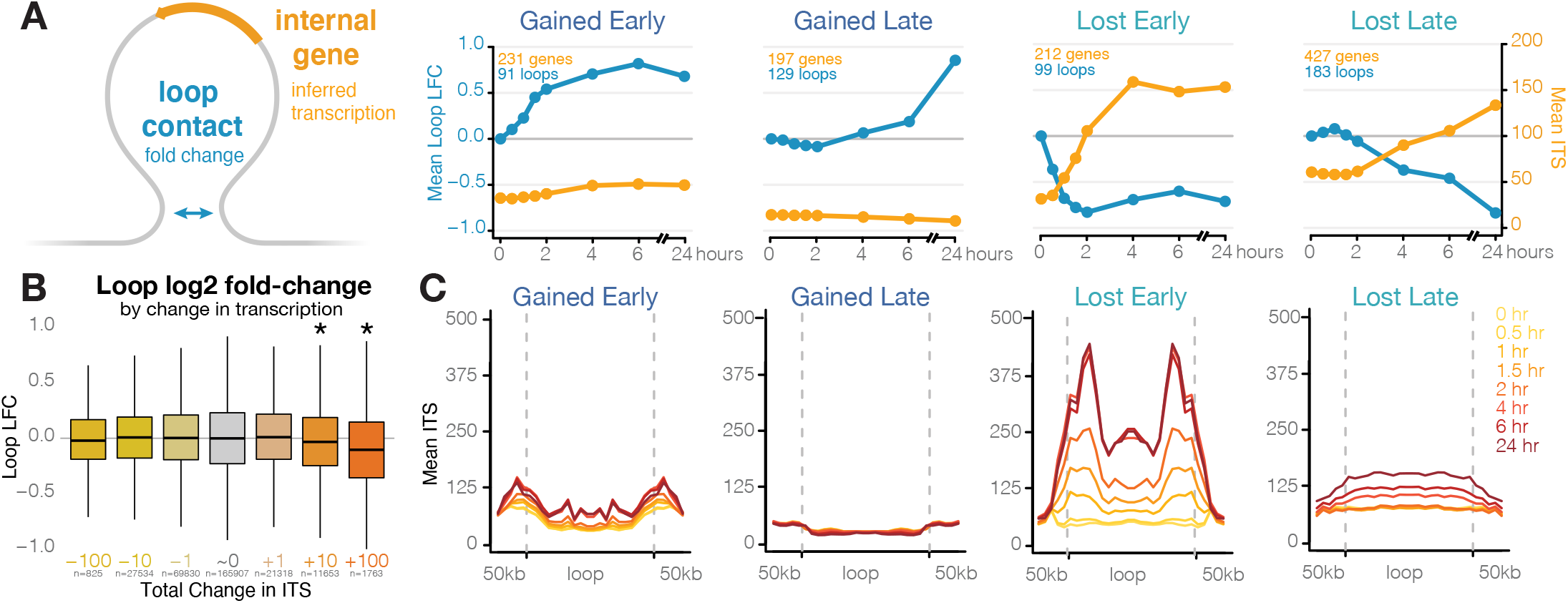
Lost loops are characterized by high levels of internal transcription. **(A)** Log2 fold-change of differential loops (blue) and average internal inferred transcription score (ITS, gold) for loops of each cluster. Gained loops have lower levels of internal transcription than lost loops and the temporal dynamics of changes in transcription are anti-correlated with changes in loop strength among lost loops. **(B)** Binning loops based on their change in internal transcription shows significant weakening of loops that gain 10 or more ITS per 10kb (Wilcoxon signed rank test, p-value p < 10-20). **(C)** Average inferred transcription score within and 50 Kb beyond loop boundaries. Transcription is highest at and beyond loop anchors in gained early loops, low among gained late loops, and localized within loop bounds in lost loops.

Taken together, this suggests that the causal arrow between looping and transcription might point both ways: DNA loop formation may contribute to increased transcription of target genes, but very high levels of transcription could weaken loops by antagonizing loop extrusion as previously observed^24,25,27,48–50^. An example of these potential phenomena can be seen at the *GBP* locus **(Fig 5A)**. In untreated cells, seven *GBP* genes are encompassed by two large (370- and 470-Kb) “structural” loops whose anchors do not overlap active gene promoters. Small loops start to form as early as 30 minutes after activation, connecting H3K27ac peaks to promoters. This is followed by increased expression of genes at the anchors of those loops. This increased expression is coupled with the loss of the large structural loops that span this locus. Visualizing these changes via line plots **(Fig 5B-E)** highlights the correlation between looping and anchor gene transcription as well as the inverse correlation between structural loops and internal transcription.

**Figure 5.**
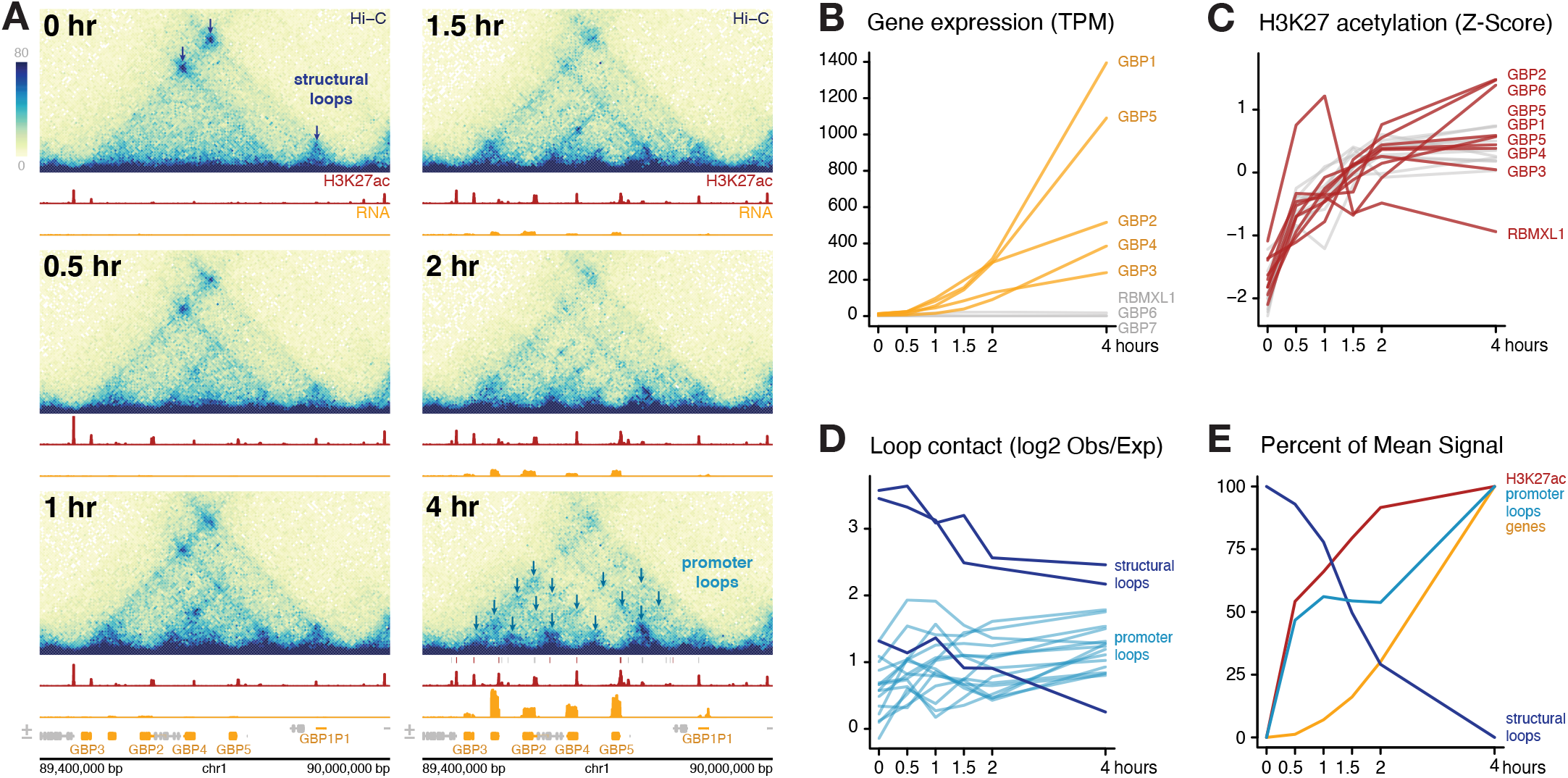
Long-distance loops are lost concurrently with increased internal transcription and restructuring at the GBP locus. **(A)** Chromatin structure, H3K27 acetylation, and gene transcription change drastically over the first 4 hours at the GBP locus of chromosome 1. Diagrams of these changes are shown on the right. Prior to treatment, two large “structural” loops (not connecting enhancers and promoters) encompass several GBP genes. After 30-60 minutes of LPS/IFNg treatment, GBP promoters become acetylated. From 1 hour onward, as acetylation increases, connections form between the GBP promoters. As genes become more highly expressed 1.5 hours and beyond, the original long-distance structural loops weaken in favor of shorter-range, transcription-correlated contacts. **(B)** The TPM of each gene within the region, with the up-regulated genes highlighted in yellow (as in figure A). **(C)** The z-score normalized change in H3K27ac at promoters (red) and putative enhancers (grey) in this region. The promoters and enhancers plotted are highlighted in the 4 hr panel of figure A. **(D)** The log-transformed ratio of observed to expected contact frequency of several points in the region. “Structural” loops (as in figure A, 0 hr) are colored dark blue, and promoter-promoter and promoter-enhancer loops (as in Figure A, 4 hr) are colored light blue. **(E)** The mean TPM (for expressed genes), z-score (for H3K27ac), and log2 observed/expected ratio (for structural or promoter contacts) for the individual features highlighted in figures B-D.

## DISCUSSION

We conducted a fine-scale multi-omic timecourse of macrophage activation and quantified changes in DNA looping, enhancer acetylation, and gene expression. The unprecedented temporal resolution of the Hi-C data revealed changes in chromatin looping along short, transcriptionally relevant timescales that were undetected at timecourse endpoints. Combining the data across timepoints yielded one of the deepest Hi-C data sets to date (over 16 billion contacts), allowing sensitive and robust detection of macrophage chromatin loops. Integration of the data revealed several novel findings regarding the nature of DNA loops.

The correlated changes we observe between looped enhancers and promoters are consistent with loops serving as a functional bridge between enhancers and their target genes, at least for genes regulated in response to external stimuli. This is further supported by temporal ordering of events that revealed that loop formation and looped-enhancer activation occur prior to increases in anchor gene expression. This agrees with a previous study of cells responding to glucocorticoids in which maximal changes in loops were observed earlier than maximal changes in genes^20^. These results are somewhat inconsistent with results from Rao *et al.* which showed very few changes in gene expression in response to global loop disruption via RAD21 degradation^21^; however, in that study the cells were grown in steady-state and were not responding to external stimuli. In an experimental setup more comparable to ours, cohesin depletion did disrupt the transcriptional response of macrophages to microbial stimuli^23^. Taken together, this suggests that loops likely do play a critical role in mediating transcriptional changes in cellular response to stimuli.

Intriguingly, we found that non-looped enhancer-promoter pairs that were matched for both contact frequency and histone H3K27ac levels did not exhibit the same level of temporal correlation as looped enhancer-promoter pairs. This suggests that loops may exert their regulatory control via mechanisms beyond merely increasing contact frequency between enhancers and promoters. One possible, albeit speculative, explanation is that activation of transcription by distal enhancers may require prolonged enhancer-promoter contact rather than overall contact frequency. Transcription factor binding is typically quite transient^51^, and prolonged contact might be required for proper formation of enhancer, polymerase, or mediator complexes that drive transcriptional activation. Recent work using 3D super-resolution live-cell imaging found that loops stabilized contact between anchors for 10-30 minutes^52^. In the absence of a chromatin loop, such prolonged contact is unlikely even for non-looped enahncers and promoters that are separated by relatively short genomic distances. Closely spaced but non-looped enhancer-promoter pairs might participate in much more frequent but shorter duration contacts that are insufficient for transcriptional activation. Hi-C data measures contact frequency but cannot differentiate between frequent short interactions and infrequent but prolonged interactions. Further exploration is required to determine if prolonged contacts do indeed account for these differences and if so, what the exact mechanisms are.

Several analyses from this paper support a model in which high levels of transcription could stall, displace, or generally antagonize loop extrusion complexes. First, we found that changes in gene expression exhibit a directional bias at differential loop anchors. The anchors of gained enhancer-promoter loops were associated with increased gene expression of genes oriented away from the center of the loop but not genes oriented towards the center of the loop. In contrast, lost enhancer-promoter loops were associated with increased expression of anchor genes oriented towards the center of the loop but not those oriented away from it. Moreover, the temporal patterns of loop loss and internal transcription were anti-correlated. These temporal analyses agree with previous work showing accumulation of cohesin at the 3’ ends of genes in a manner correlated with the amount of transcription and also sensitive to transcription inhibition^24,27,28^, and recent studies demonstrating that RNA polymerase may act as a “moving barrier” to loop extrusion^49^. Transcription may also shape chromatin independently of cohesin. Recent high-resolution microscopy and Micro-C experiments have detected fine-scale cohesin-independent structures between and within highly expressed genes, which could compete with or disrupt cohesin-mediated structures^50,53^. Transcription inhibition interrupts these local structures but leaves intact broader loops, domains, and compartments. Additionally, virtually all transcriptionally active chromatin exhibits elevated contact frequency via a phenomenon called compartmentalization that does not require cohesin^21,47,54^. It remains possible that such compartmentalization itself could disrupt loops surrounding highly expressed genes.

Finally, lost loops were associated with relatively high levels of internal transcription and only very large changes in transcription were associated with decreased looping. This might reflect the fact that, in most cases, collisions between transcription and loop extrusion are rare. Indeed, transcription occurs in relatively infrequent bursts^55^, and loops appear to spend at least some time in either fully looped or fully non-looped states^52^. So at low to moderate levels of transcription, collisions might be uncommon and are not a major driver of 3D chromatin structure. So perhaps, it is only at extremely high levels of transcription where such collisions are frequent enough to lead to observable losses in loop-based contacts. Alternatively, it is possible that transcription only slightly impedes extrusion, and at low levels of transcription, changes in contact frequency are imperceptible. This agrees with studies showing that transcription briefly stalls condensin translocation but that it only measurably impacts 3D chromatin structures at extremely highly expressed loci such as at rRNA genes^25^.

This fine-scale timecourse of looping in human macrophages provides insight into the temporal organization of regulatory events in human cells responding to external stimuli and a deeper understanding of the mechanisms driving transcriptional regulation in human cells. Some of the findings could be useful for predicting functional enhancer-promoter pairs. For example, the temporally coordinated changes observed at looped enhancer-promoter pairs could be employed to refine and potentially improve predictions made by the activity-by-contact model^56^.

This work supports a model in which loop extrusion and transcription participate in a coordinated dance and can influence each other in a reciprocal relationship. If this holds true, it could have important implications for how genes are organized within the context of chromatin loops. For example, genes oriented towards the center of a loop could be regulated by a negative feedback mechanism in which high levels of transcription might decrease looping between the promoter and a distal enhancer. Moving forward, incorporation of more data types into these timecourses should reveal further insights into the mechanisms of 3D chromatin structure and gene regulation.

## METHODS

### Macrophage differentiation and activation

THP-1 monocytes were grown and maintained in RPMI media with 10% fetal bovine serum (FBS) and 1% penicil-lin-streptomycin (PS).

For differentiation into macrophages, monocytes were transferred to 6-well plates (RNA-seq, ATAC-seq) or T-175 flasks (Hi-C, ChIP-seq) at a density of 2×10^5^ cells/mL and treated with 25 nM PMA for 24 hours, over which time the cells become adherent. The media was then aspirated off, the flasks were washed gently with RPMI, and then fresh RPMI (10% FBS, 1% PS) and rested for 72 hours.

The resting macrophages were then treated with a combination of 10 ng/mL lipopolysaccharide (LPS) and 20 ng/mL interferon gamma (IFNγ) in fresh RPMI (10% FBS, 1% PS). Cells were harvested without treatment, or 0.5, 1, 1.5, 2, 4, 6, or 24 hours after LPS and IFNγ treatment.

During each treatment, extra 0- and 2-hour samples were prepared simultaneously for RNA extraction, and qPCR was used to measure the regulation of FOS, IL1B and IL6 to confirm consistent treatment response.

For all library preparations, the differentiation and activation treatment was performed from freshly thawed THP-1 cells on two separate occasions, to achieve the closest approximation to two biological replicates using cultured cell types.

### Crosslinking

For ChIP-seq and Hi-C, cells were grown in T-175 flasks, each containing 20×10^6^ cells at a density of 2×10^5^ cells/ mL. Cells were crosslinked using 1% formaldehyde in RPMI for 10 minutes with gentle shaking. Crosslinking was then quenched with 10% 2.0 M cold glycine for 5 minutes. The media was then removed and cells were scraped into cold PBS. Each flask was divided into 4 tubes of approximately 5×10^6^ cells each. Cells were spun down at 526 G for 5m, resuspended in PBS and respun to wash away residual formaldehyde. Cells were then frozen in liquid nitrogen and stored at −80 for library preparation.

### RNA-seq library preparation

RNA was extracted using the QIAGEN RNeasy Mini kit with DNase I treatment. RNA integrity numbers were confirmed using a Tapestation RNA screentape to be above 9.8, and a Qubit High Sensitivity assay was used to determine RNA concentration.

Ribosomal RNA was removed using the NEB rRNA Depletion Kit (Human/Mouse/Rat) using 500 ng of RNA as input. Following depletion, RNA-seq libraries were prepared using the NEB Ultra II Directional RNA Library Prep Kit for Illumina, and NEBNext Multiplex Oligos for Illumina. Library concentration and fragment size was determined using Qubit (dsDNA HS assay) and Tapestation (D1000 screentape). Libraries from each timepoint were pooled to a final DNA concentration of 15 nM, and 75-bp paired-end reads were sequenced on an Illumina NextSeq 500 using a High Output Kit.

### ChIP-seq library preparation

Four frozen cell pellets (5×10^6^ cells each) were used for each timepoint. Pellets were first rinsed in 10 mL rinse buffer 1 (50 mM HEPES pH 8, 140 mM NaCl, 1 mM EDTA, 10% glycerol, 0.5% NP-40, Triton-X), incubated on ice for 10 minutes, and then spun down at 2,400 G at 4°C for 5 minutes. Supernatant was removed and the pellets were rinsed again in rinse buffer 2 (10 mM Tris pH 8, 1 mM EDTA, 0.5 mM EGTA, 200 mM NaCl), and spun at the same settings. Supernatant was removed, and 5 mL of shearing buffer (10 mM Tris pH 8, 2% Triton-X, 1% SDS, 100 mM NaCl, 1 mM EDTA) was added to the tubes to wash out the rinse buffer. The samples were centrifuged at 2,400G at 4°C for 3 minutes, the shearing buffer was removed and this step was repeated. The cell pellets were then resuspended in 88 uL of shearing buffer, 2 uL of protease inhibitor cocktail (PIC), and 10 uL of nanodroplets (Triangle Biotechnology, Inc.) per 10 million cells^57^. Samples were aliquoted into 100 uL tubes and sheared using a Covaris E110 (intensity 6, 210 seconds). Cells were spun down at max speed for 2 minutes and the supernatant was retained.

In order to determine the concentration of chromatin, 10 uL was removed (while the rest of the sample was stored at −80°C), and crosslinking was reversed by adding 5 uL of 5M NaCl, 125 uL of TE buffer (10 mM Tris pH 8, 1 mM EDTA) and 125 uL of elution buffer (1M Tris pH 8, 10 mM EDTA, 1% SDS), vortexed, and incubated overnight at 65°C. Samples were spun down and added 7.5 uL of proteinase K and 3 uL of RNase A. DNA was extracted using the Zymo ChIP DNA Clean & Concentrator Kit, quantified using Qubit (dsDNA broad-range (BR) assay), and run on a gel to ensure fragment sizes of 100-300 bp and concentrations high enough to continue with library prep.

Immunoprecipitation of the remaining volume from each sheared sample was completed using the Active Motif ChIP-IT High Sensitivity kit, using 2.8 ug of chromatin from each timepoint (as determined by the lowest yield samples), and 4 ug of anti-H3K27ac antibody (AbCam ab4729). Following overnight antibody incubation and washing steps, crosslinking was reversed by adding 100 uL of elution buffer (described previously) and 4 uL of 5M NaCl to 100 uL of the IP reactions, vortexing, and then incubating overnight at 65°C. DNA was purified using the Zymo ChIP DNA Clean & Concentrator kit, and quantified using Qubit (dsDNA high-sensitivity (HS) assay), as before.

Following the final dilution, libraries were prepared using the NEB Ultra II DNA Library Prep Kit with NEBNext Multiplex Oligos for Illumina with 0.88 ng of DNA as input. Libraries were analyzed using Qubit (dsDNA HS assay) and Tapestation (D1000 screentape) and pooled to a final concentration of 12 nM, and then 75-bp paired-end reads were sequenced on an Illumina NextSeq 500 using a High Output Kit.

### *In Situ* Hi-C Library Preparation

Three treatments (biological replicates) were conducted, and one or two frozen cell pellets (5×10^6^ cells each) were used to generate separate libraries as technical replicates (1 technical replicate for first biological replicate; 2 technical replicates for second and third biological replicates). Libraries were prepared using the in situ Hi-C protocol as described in Rao *et al.* 2014^9^. In brief, cross-linked cells were lysed on ice, nuclei were isolated, and chromatin was digested overnight with the MboI restriction enzyme. Chromatin ends were biotinylated, proximity ligated, and crosslinking was reversed. Samples were sheared on a Covaris LE 220 (DF 25, PIP 500, 200 cycles/ burst, 90 seconds), quantified using Qubit (dsDNA High Sensitivity (HS) assay), and a small sample was run on an agarose gel to ensure proper fragmentation. DNA sized 300-500 bp was selected for using AMPure XP beads, and then eluted. Biotinylated chromatin was then pulled down using streptavidin beads. Following removal of biotin from unligated ends and repair of sheared DNA ends, unique Illumina TruSeq Nano (Set A) indices were ligated onto the samples. Libraries were amplified off of streptavidin beads using 7-10 PCR cycles based on post-size selection concentrations, quantified again using a Qubit (dsDNA HS assay), and fragment length was determined using Tapestation (D1000 screentape). Libraries were pooled to 10 nM. Paired-end 150-bp reads were sequenced on one or two lanes of an Illumina NovaSeq S4.

### ATAC-seq Library Preparation

ATAC-seq libraries were prepared using the Omni ATAC-seq protocol as described in Corces *et al.* 2017^58^. Adherent macrophages were washed once with PBS and lifted off of the plate with EDTA for 5 minutes. EDTA was quenched with RPMI, and library preparation was performed on 50,000 cells. 3.75 μL of Illumina Nextera XT indices were used in PCR and qPCR.

After performing the initial 5 cycles of PCR, 5% of the PCR reaction was used in qPCR to determine how many additional cycles were required. 4-7 cycles were determined to be sufficient for the final amplification. A 2-sided bead cleanup with AMPure XP beads was performed (0.5X, then 1.3X). Libraries were quantified using Qubit (dsDNA HS Assay) and the KAPA Library Quantification kit. Libraries from each time-point were pooled to a concentration of 8 nM or 10 nM for each biological replicate, and 75-bp paired-end reads were sequenced on an Illumina NextSeq 500 using a High Output Kit.

### RNA-seq processing and gene quantification

Adaptors and low-quality reads were trimmed from paired-end reads using Trim Galore! (version 0.4.3). Salmon (version 1.4.0) was used in quant mode to quantify reads to hg19 transcripts from GENCODE (version 19)^59,60^. For signal tracks, reads were aligned using HISAT2 (version 2.1.0), indexed and replicates were merged with samtools (version 1.9), and converted to bigwigs using deeptools (version 3.0.1)^61–63^. Reads were summarized to a gene level using tximport (R version 3.3.1, tximport version 1.2.0), which was then used as input for differential analysis in DESeq2 (version 1.33.5)^43,64^. FastQC and MultiQC were used to assess library quality metrics (version 0.11.5, 1.5)^65,66^.

### ATAC- and ChIP-seq processing and peak calling

Adaptors and low-quality reads were trimmed from paired-end reads using Trim Galore! (version 0.4.3)^59^. Reads were aligned using BWA mem (version 0.7.17) and sorted using Samtools (version 1.9)^62,67^. Duplicates were removed with PicardTools (version 2.10.3) and for ATAC-seq libraries, mitochondrial reads were removed using Samtools idxstats^68^. Samtools was also used to merge replicates for each timepoint, and index BAM files. Peaks were called from the merged alignments using MACS2 with the following settings: -f BAM -q 0.01 -g hs --nomodel --extsize 200 --keep-dup all -B --SPMR (version 2.1.1.20160309)^69^. ChIP-seq peaks used the MACS2 setting --shift 0, while ATAC-seq peaks used --shift 100. Peaks from all timepoints were then merged using bedtools (version 2.28), generating 118,344 ChIP-seq and 193,853 ATAC-seq peaks in total. For each replicate BAM file, ChIP-seq counts were extracted from ATAC-seq peak locations using bedtools multicov^70^. Bedtools intersect was used to subset for ATAC-seq peaks that overlapped H3K27ac ChIP-seq peaks, and these 89,503 peaks were considered putative regulatory regions. Raw counts at these enhancers (8 timepoints, 2 replicates each) were used as input for differential analysis with DESeq2 (version 1.33.5)^43^. Signal tracks were made from alignments using deeptools (version 3.0.1)^63^.

### Enhancer and promoter definitions

Gene promoters were identified as regions 2,000 base pairs upstream and 200 base pairs downstream of gene transcriptional start sites (TSS). Promoter H3K27ac signal was calculated based on any overlapping H3K27ac and ATAC-seq peaks within promoter regions using bedtools intersect (version 2.28)^70^. Enhancers were identified as overlapping H3K27ac and ATAC-seq peaks that did not overlap with defined promoter regions.

### Hi-C processing, loop and compartment calling

Hi-C data was processed using the Juicer pipeline as initially described in Rao *et al.* (version 1.5.6)^9^. Hi-C maps were made at 5 and 10 kb resolution for each technical replicate (8 timepoints, each with 5 technical replicates across 3 biological replicates), as well as for each timepoint (all replicates combined). Additionally, a “Mega” map from all timepoints was made, also using Juicer.

Loops were identified at 5 kb using SIP (version 1.6.1)^42^. Loops were called from the individual timepoint maps using the settings “-g 2 -t 2000 -fdr 0.05”, and from the Mega map with the settings “-g 1 -t 2000 -fdr 0.05”. The loops were then extrapolated to 10kb, concatenated, and merged in R using DBScan (version 1.1.8) with an epsilon of 20 kb (manhattan distance), keeping the mean of modes for coordinates, resulting in 42,690 total loops. The counts for these loops were then extracted from the Hi-C files of each technical replicate (un-normalized, 10kb resolution) using strawr (version 0.0.9)^71^. These raw counts (8 timepoints, 3 biological replicates each, two with 2 technical replicates and one with 1 technical replicate each) were used as input for differential analysis with DESeq2^43^.

### Differential gene and peak analysis

DESeq2 was used for differential analysis of genes, loops, and peaks^43^. Each analysis used a likelihood ratio test (LRT), with a full design of “~bioRep + time” and a reduced design of “~bioRep”. Posterior log-2 fold changes (LFC) were estimated using apeglm^72^. Significant results were determined based on an absolute LFC greater than 1 and an adjusted p-value below 0.01.

Raw counts were converted into Z-scores by first conducting a variance-stabilizing transformation across all features, and then centering and scaling the data in each feature based on standard deviations from the mean. Genes were categorized into up- and downregulated based on the signage of their Z-score at 0 hours of LPS/ IFNg treatment, and then sorted based on their timepoint of maximum Z-score.

### Differential loop analysis and clustering

DESeq2 was also used for differential analysis of loops^43^. Differential analysis used a likelihood ratio test (LRT), with a full design of “~techRep + bioRep + time”, and a reduced design of “~techRep + bioRep”. Significant results were determined based on an absolute LFC greater than 0.585 (fold-change of ±1.5) and an adjusted p-value below 0.05.

Raw counts were converted into Z-scores by first conducting a variance-stabilizing transformation across all features, and then centering and scaling the data in each feature based on standard deviations from the mean. These Z-scores were then used to cluster loops using k-means clustering (k=4). For the survey of loop contacts at the GBP locus, log2 observed/expected KR-normalized counts were extracted using strawr.

### Matched enhancer-promoter sets

Covariate-matched subset selection among non-looped enhancer-promoter pairs was performed using the matchRanges function from the nullranges package. Enhancer-promoter pair distance or total contact frequency were used as covariates. Total contact frequency was calculated from KR normalized counts from the combined Mega map, effectively a sum of contacts across all timepoints and replicates. Matching was done with the stratified matching method without replacement. Enhancer strength, defined by the sum of H3K27ac variance-stabilized counts across all timepoints and replicates, was compared between the looped and matched non-looped sets.

### Inferred transcription score (ITS) calculations

Inferred transcription scores (ITS) were calculated in order to estimate the degree of transcription occurring throughout gene bodies, including introns, as extrapolated from the mature mRNA TPM levels. Gene-level TPM as quantified by Salmon and summarized by txImport (see RNA-seq processing methods). The genome was binned into 10kb regions using bedtools makewindows (version 2.28), and then overlapped with gene bodies^70^. Gene TPM values were applied to each overlapping bin, adjusted based on the percentage of bin overlap. For example, a gene of TPM 50 with a TSS at position 100,000 bp and a TTS at position 115,000 bp would contribute an ITS of 50 to the bin of 100,000-110,000, and an ITS of 25 to the bin of 110,000-120,000. In bins with multiple genes, ITS scores were generated by summing the TPM contribution from each gene.

## Data availability

Raw and processed data for Hi-C (GSE201353), RNA-seq (GSE201354), ATAC-seq (GSE201351), and ChIP-seq (GSE201352) data are publicly available on GEO and SRA under SuperSeries GSE201376.

## ACKNOWLEDGEMENTS

We thank Erika Deoudes for data visualization, illustration, proof reading, and typesetting. We thank Samantha Pattenden for use of the Covaris LE220 instrument which was provided by the North Carolina Biotechnology Center Institute Development Program grant 2017-IDG-1005. We thank Samantha Pattenden and Paul Dayton for generously supplying the nanodroplets used for chromatin shearing (Triangle Biotechnology FF101-5000, Durham, NC). This work was supported by NIH grants (R35-GM128645 to D.H.P.; R00HG008662 to D.H.P.; R35GM143532 to I.H.; R01AG066871 to H. W.) and multiple NIH training grants (T32-GM067553 for E.S.D, T32 GM007092 for K.S.M.R. and E.A.T., T32 GM135128 for M.L.B.). I. Y.Q. was supported by a BrightFocus Foundation postdoctoral fellowship. I.H. was supported by an award from the Cancer Prevention & Research Institute of Texas (RR170030). E.S.D. and M.I.L were supported by a CZI Essential Open Source Software for Science (EOSS) Round 3 award.

## AUTHOR CONTRIBUTIONS

K.S.M.R. designed and performed the majority of experiments, performed computational analysis, and wrote the paper.

E.S.D. developed software and performed computational analysis M.L.B. performed some cell culture and genomic library preparation experiments

A.C. performed some cell culture experiments

E.A.T. performed some cell culture experiments and assisted with ChIP-seq

I.Y.Q. prepared ATAC-seq libraries

S.C. helped prepared ATAC-seq libraries and cross-linking for ChIP-seq

K.W. assisted in cell culture experiments

H.W. supervised and help interpret genomic data

I.H. oversaw the planning, supervising, and interpretation of some experiments

M.I.L. supervised computational analysis and software development

D.H.P. acquired funding, conceptualized the project, supervised experiments and data analysis, and helped write the paper.

## SUPPLEMENTAL FIGURES

**Figure S1.**
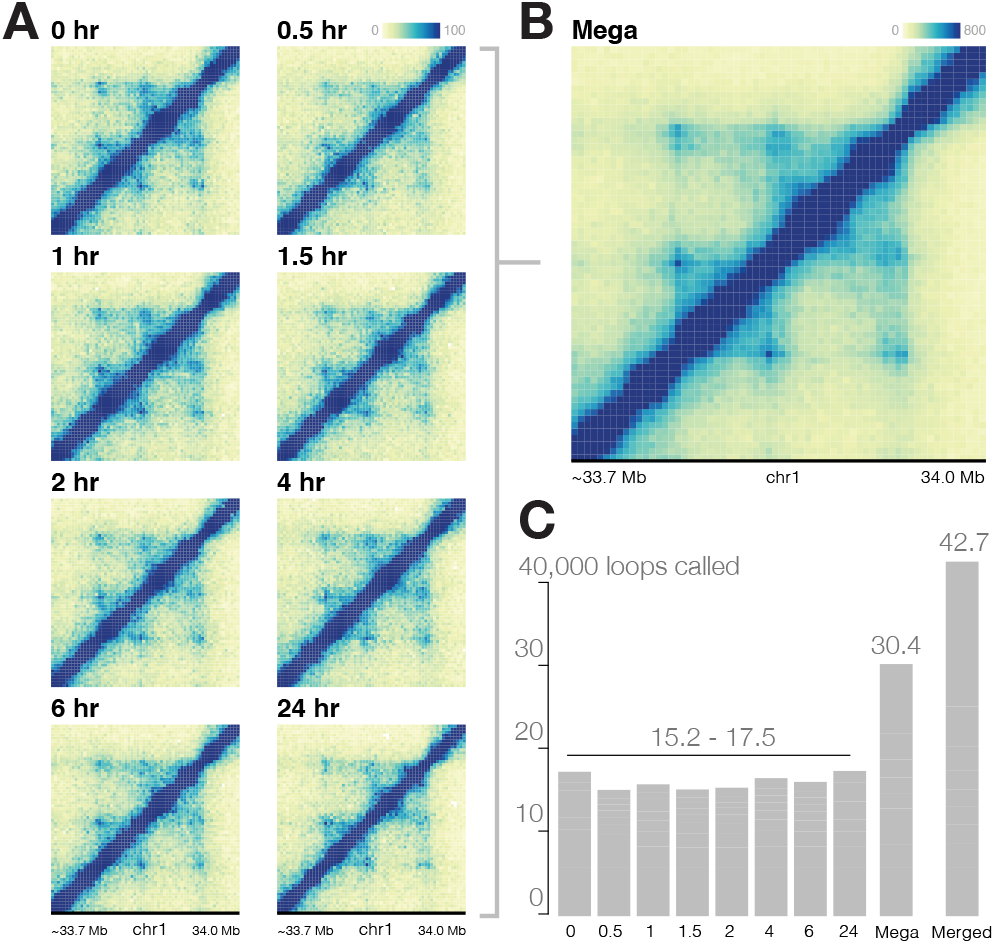
Deeply sequenced in situ Hi-C sensitively identifies loops. **(A)** Hi-C maps of roughly 2 billion Hi-C contacts were generated for each timepoint. **(B)** For added depth and sensitivity of loop detection, each timepoint was merged into a Mega map of 15.7 billion Hi-C contacts. **(C)** Loops were called in each individual timepoint, as well as the Mega map, and loops with both anchors within 20 kb were merged. Nearly twice as many loops were called from the Mega map compared to individual timepoint maps, resulting in 42,690 total chromatin loops.

**Figure S2.**
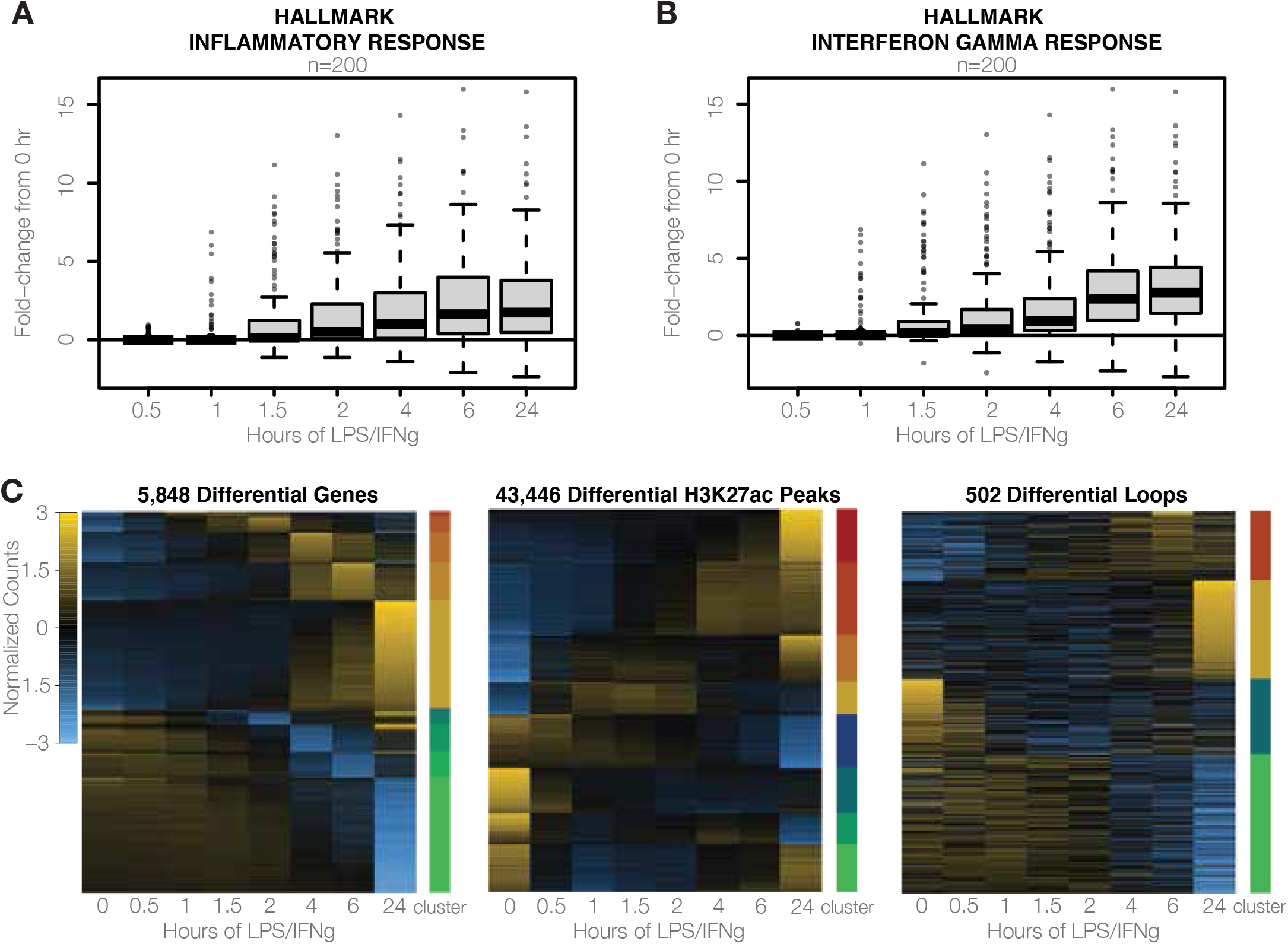
Transcriptional profile consistent with inflammatory hallmarks. **(A)** Genes previously identified as part of the hallmark inflammatory response and **(B)** hallmark interferon gamma response were investigated in this system. Canonically upregulated genes exhibit positive fold-change in response to LPS/IFNg, especially at 4 hours and beyond. **(C)** Differential genes, overlapping ATAC-seq and H3K27ac ChIP-seq peaks, and loops occur at multiple timescales.

**Figure S3.**
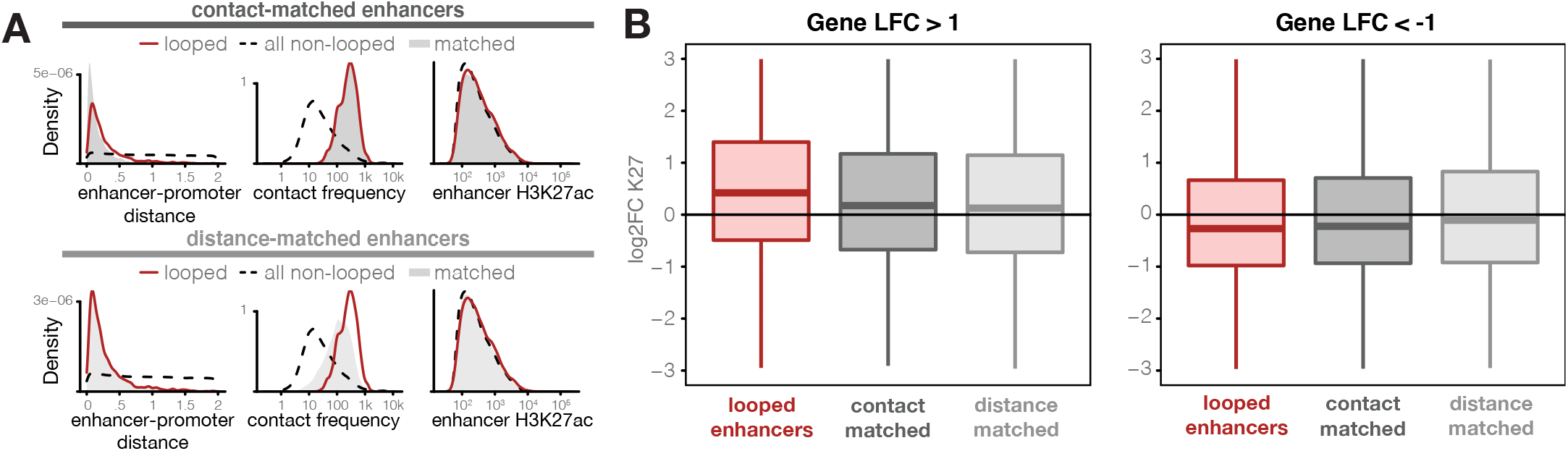
Looped enhancer-promoter pairs correlate in other systems. **(A)** Contact- and distance-matched enhancer-promoter pairs were identified to compare against looped enhancer-promoter pairs seen in monocyte-to-macrophage differentiation. **(B)** Enhancers looped to upregulated genes show a correlated increase in H3K27ac that is significantly higher than both contact- and distance-matched pairs (Wilcoxon rank-sum test, p-value < 10-5). Similarly, enhancers looped to downregulated genes show a correlated decrease in H3K27ac that is significantly higher than distance-matched pairs (Wilcoxon rank-sum test, p-value < 10-5).

**Figure S4.**
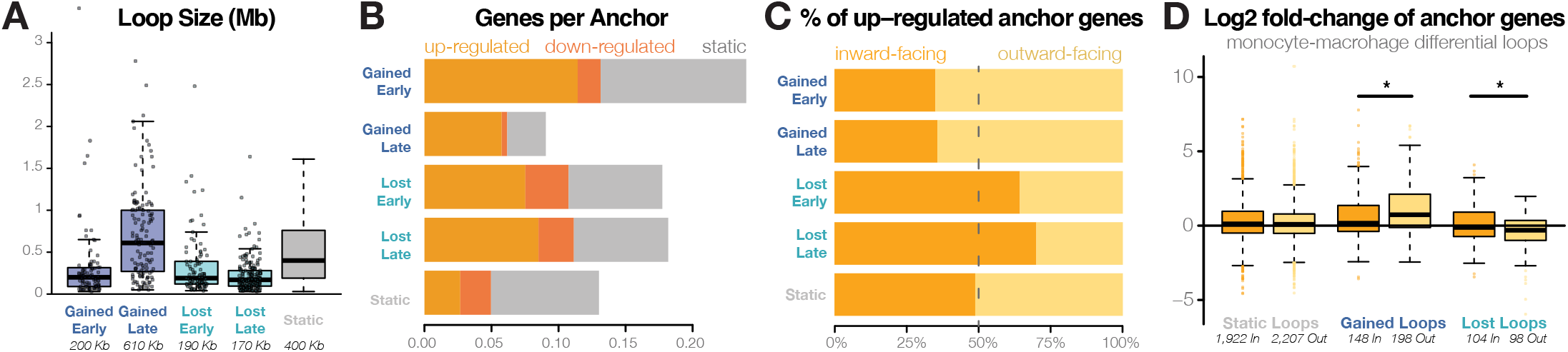
Differential loop features. **(A)** The distribution of loop sizes based on differential cluster. **(B)** The average number of genes per loop anchor for loops of each differential cluster, distinguished by differential status of anchor genes. **(C)** The percentage of upregulated anchor genes which are inward - or outward-facing among each class of differential loops. **(D)** Distributions of log2 fold-changes of genes with promoters in the anchors of static and differential loops from monocyte-macrophage differentiation^19^. At gained loop anchors, the fold-change of outward-facing genes is significantly higher than inward-facing genes, while the opposite trend is seen among genes at lost loops (Wilcoxon rank-sum test, p-value < 0.05).

